# Multilayer Label Free Non-Faradic Electrochemical Impedance Immunosensor for Cortisol Detection

**DOI:** 10.1101/2023.07.21.550009

**Authors:** Chinmay Gupta, Sudip Kumar Pattanayek, Biswarup Mukherjee, Sachin Kumar

## Abstract

Cortisol, a well-known psychological stress biomarker, produced by the hypothalamic-pituitary-adrenal system, tends to intensify with stressors. Prolonged overexpression of cortisol leads to chronic stress that causes disparities in the proper functioning of the human body. Thus, there is a huge demand for developing a rapid cortisol detection system. Several point-of-care diagnostic techniques are available for rapid cortisol detection, such as electrochemical sensing, which works on changes in the electrical properties due to the binding of an analyte with a biorecognition element. Researchers have used different electrochemical methodologies such as cyclic voltammetry (CV), chronoamperometry, and faradic electrochemical impedance spectroscopy (EIS) for the detection of cortisol, but usage of external redox active reagents, low sensitivity, limited dynamic range, and electrode fouling nature limits their use. Hence, we reported a label-free and non-invasive cortisol detection using non-faradic EIS. A novel multilayer immunosensor was fabricated on PEDOT: PSS coated ITO glass by functionalizing with cortisol antibodies. Specific and rapid detection of cortisol was measured by monitoring the change in impedance in a dynamic range from 50-200 ng/mL. We envision the developed immunosensor has the potential for new developments in stress monitoring, disease prognosis, and enable personalized care.

**Highlights:**

- Novel PEDOT: PSS based multilayer immunosensor for cortisol detection
- Impedance based label free detection of cortisol using non-faradic EIS
- Presentation of detailed multilayer immunosensor fabrication, experimental detection, and equivalent circuit model with working mechanism
- Cortisol detection in a dynamic range of 50-200 ng/mL

**Graphical Abstract:** 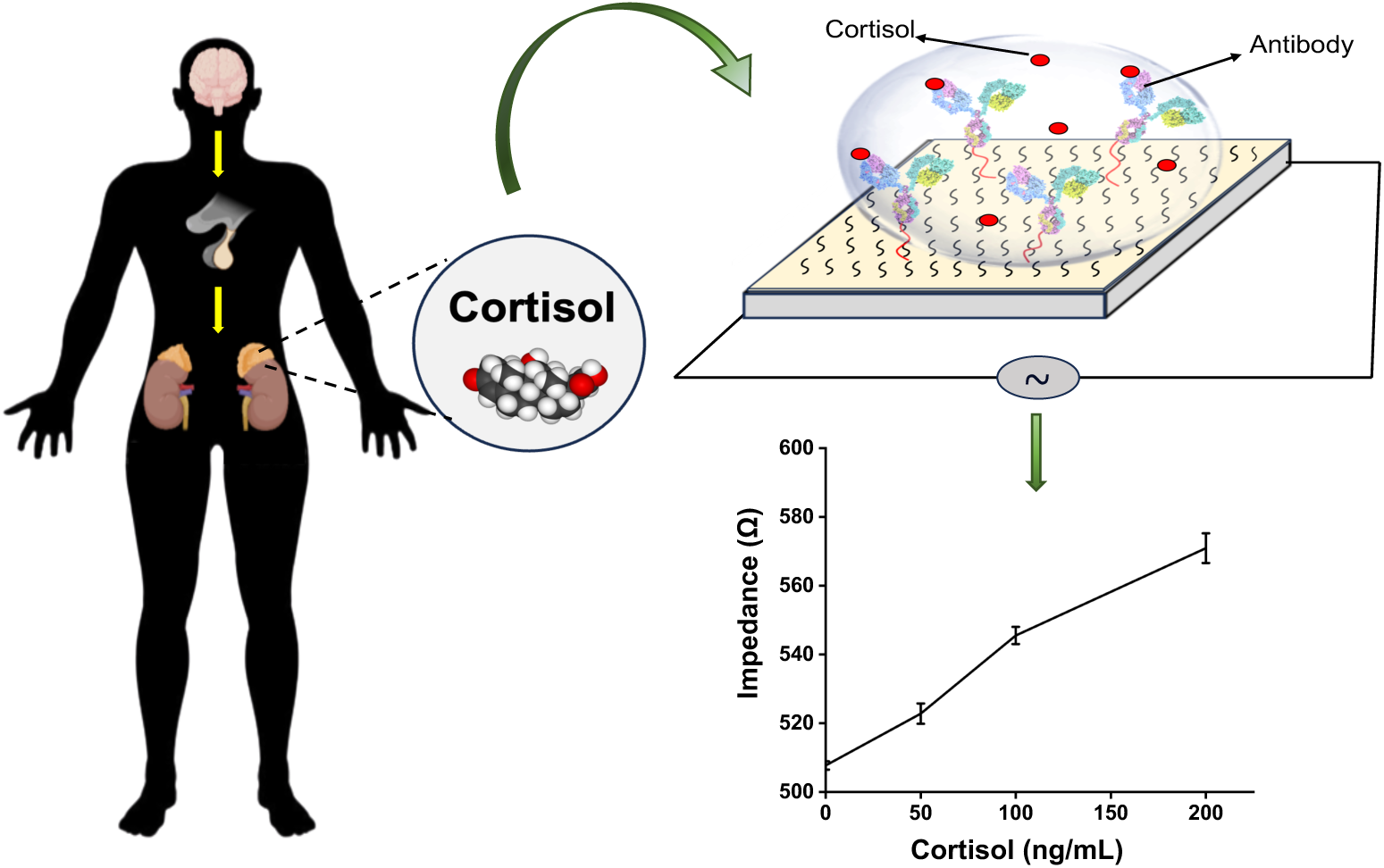

## 1. Introduction

Rising global psychological stress caused by fast-paced modern lifestyle activity linked with an increasing risk of numerous health problems, comprising autoimmune disorders, cardiovascular diseases, and mental illnesses [1]. One crucial mechanism connecting between psychological stress and health problems is a hormone called cortisol, produced by the hypothalamic-pituitary-adrenal (HPA) axis as a result of body’s response to stress and thus called “stress hormone”[2–4]. As a stress hormone, cortisol controls several physiological functions such as heart rate, blood glucose level, immune response, metabolism and central nervous system activation [5,6]. In response to psychological and physical stress, the body releases cortisol (in blood serum, saliva, and sweat) in a pulsatile pattern to maintain homeostasis. Under normal conditions, cortisol exhibits a distinct circadian rhythm, with its highest concentration in the morning (50-250 ng/mL) and lowest around midnight (30-130 ng/mL) in serum [4,7]. Moreover, in response to sudden physical stress like injury, intense exercise, imminent risk, and perceived threat leads to rapid release of cortisol and enables the body to adapt to stressful environments and restore homeostasis [8,9]. However, prolonged psychological stress leads to the persistent increase in cortisol level and has devastating effects on different body systems like cardiovascular diseases, mental disorders, and a compromised immune system [8,10,11]. Hence, assessing cortisol level becomes a vital biomarker for stress monitoring in normal as well as diseased patients.

Several conventional methods like radioimmunoassay (RIA), enzyme-linked immunosorbent assay (ELISA), high-performance liquid chromatography (HPLC), electrochemical-luminescent immunoassay (ELICA), etc.) has been developed for detection of cortisol in different biological fluids such as saliva, sweat, urine, blood serum, etc. [12–16]. Although these conventional methods provide a snapshot of the cortisol level in biological fluids, but they are labor-intensive, chronophagous, require multistep sample preparation, and rely significantly on high end equipment and skillful technician. With advance technological developments, researchers are transitioning from traditional medical inspection to advance health-monitoring systems. Several efforts have been made to develop advanced point-of-care diagnostic devices based on quartz crystal balance [16], surface plasmon resonance [17], and electrochemical methods [18] to accurately measure cortisol level in a rapid, continuous, and non-invasive manner.

Electrochemical immunosensing is emerged as the most promising tool in advanced point-of-care diagnostics, due to its miniaturization, ease of fabrication, and continuous monitoring of cortisol [19–21]. Electrochemical immunosensing operates on the principle of detecting the change in the electrical properties of a conductive material. It is achieved through the interaction or adsorption of an analyte with antibodies on the sensor surface [22]. In recent years, various electrochemical sensing techniques have been employed to detect cortisol, including chronoamperometry [23],cyclic voltammetry (CV) [18], and faradic electrochemical impedance spectroscopy (EIS) [24]. Although all these electrochemical techniques offer appealing prospects for sensing cortisol hormone, but require an external redox active species (potassium ferro/ferricyanide), as cortisol doesn’t have active redox center [4,25]. Currently, researchers are exploring non-faradic EIS as a potential alternative to faradic EIS. It eliminates the need of chemical changes or redox active species at the electrode surface. Instead, it relies on the change in the capacitive and the resistive comportment of the sensor, resulting from explicit binding of analyte with analogous antibodies [26,27]. Using non-Faradic EIS, Arya et al. developed a capacitive aptasensor for breast cancer detection. They fabricated a gold electrode conjugated with a thiol-terminated DNA aptamer to detect human epithelial growth factor receptor 2 (HER2). The Non-Faradic EIS measurements were utilized to evaluate biosensor performance via monitoring capacitive change. The binding of HER2 on the electrode sensor showed the detection of HER2 in a range from 1 pM to 100 nM in both buffer and undiluted serum [28].

On a similar front, cortisol expression in response to physiological stress by HPA axis (**Fig. 1(i))** in multiple biofluids, including (saliva, sweat, blood and urine) **(Fig. 1(ii))** unleashes the potential for non-invasive detection and offers innovative approaches to stress management shown in **Fig. 1(iii)**. In this work, we developed a novel label-free multilayer immunosensor to measure cortisol in a buffer solution. The multilayer sensor was fabricated on indium-tin-oxide (ITO) coated glass substrate by spin coating a highly conductive poly(3,4-ethylene-dioxythiophene) polystyrene sulfonate) (PEDOT: PSS) film and silanized to link 3-aminopropyl-triethoxysilane (APTES) for conjugation of antibodies against cortisol using EDC-NHS coupling [31]. The electrochemical response of the multilayer immunosensor was acquired using Bode 100 analyzer by measuring the change in resistive behaviour of the sensor after specific binding with different cortisol concentrations. Our developed non-faradic EIS immunosensor showed specific dosage-dependent cortisol detection, which can be easily miniature into a small point-of-care device for rapid quantification of biomarkers for healthcare applications.

**Fig. 1:**
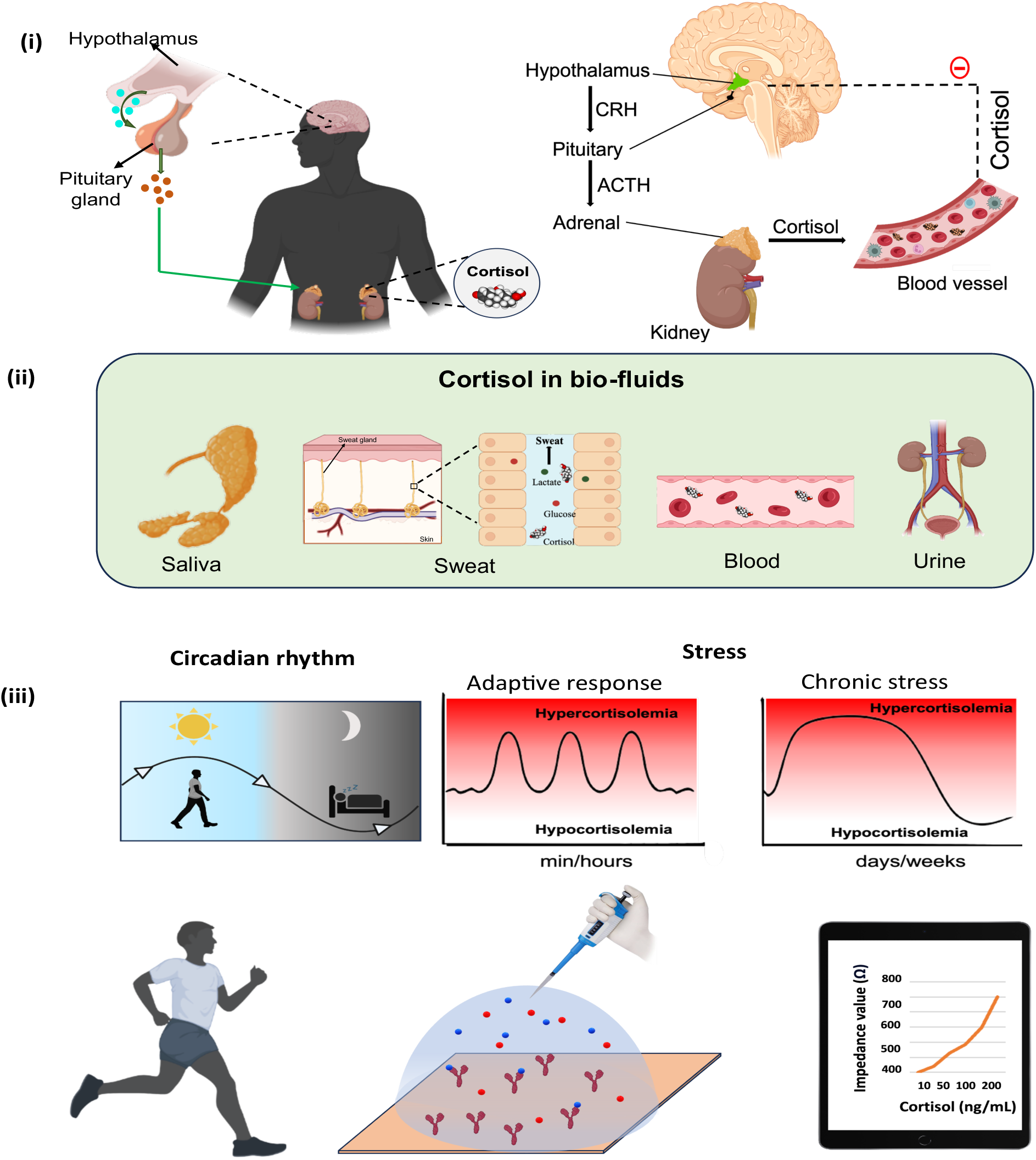
Cortisol release pathway, its partitioning in biofluids, and pulsative pattern of cortisol levels. (i) The HPA (hypothalamus-pituitary-adrenal) axis controls the release of cortisol in response to circadian rhythm and stress, (ii) Distribution of cortisol in different biofluids (saliva, sweat, blood, and urine), (iii) Fluctuation in cortisol release in response to circadian rhythm and stress. Overview of multilayer sensor and non-faradic EIS response.

## 2. Materials and Methods

### 2.1. Cortisol immunosensor fabrication

For the development of a two-electrode system, obtained ITO glass (Techinstro) was first thoroughly cleaned in following steps. First, it was immersed in a 1% surfactant solution (Labolene) and subsequently sonicated for 15 mins in different solutions of acetone, ethanol, and isopropyl alcohol (IPA) respectively. Following sonication, the glass was dried by gently purging of nitrogen gas. A UV curable resin (Creality) was manually deposited on the ITO-coated glass surface to create two masked areas for the electrodes on either ends, leaving the center unmasked and cured under a UV source for 2 min. The masked slide was then etched with a concentrated HCL (35%) solution for 45 sec to remove the ITO from the unmasked layers. The UV-cured resin was cleaned using an IPA solution. The etched glass was treated in an ultraviolet ozone (UVO) chamber for 20 min to make hydrophilic surface for the coating of conductive PEDOT: PSS (Sigma Aldrich, cat. #:65520) film. Immediately after UVO treatment, 250 µL of PEDOT: PSS (3-4% in H2O) solution was drop casted on the ITO glass substrate and spin-coated (NT12000, Navson technologies) at 1200 rpm with an acceleration of 600 rpm for 30 sec. The spin-coated samples were vacuum dried at room temperature for 24 h. Following drying process, PEDOT: PSS thin film underwent a treatment with UV-Ozone (PSPD-UVT, Novascan) to introduce polar functional groups on PEDOT: PSS thin film surface. After UV-Ozone-treatment, PEDOT: PSS film was immediately submerged in 3% APTES (SRL, cat. #:33993) in an anhydrous toluene solution to impart an amine group on PEDOT: PSS film surface that helps in conjugation antibody [29].

### 2.2. Cortisol antibody functionalization on sensor surface

By utilizing the free amine group provided by APTES, cortisol antibody was immobilized on the sensor surface using classical EDC-NHS chemistry [30,31]. In brief, EDC (Sigma Aldrich, cat. #:80097) and NHS (97%, SRL, cat. #47913) were dissolved in 0.1M MES (99%, SRL, cat. #69408) at pH 6 with concentrations of 0.2 M and 0.6 M respectively. The EDC-NHS mixture was treated with cortisol antibody (Sigma Aldrich, cat. #MAB1243) at a 500 μg/mL concentration. Antibody-treated EDC-NHS solution (30 μL) was added to the sensor surface and allowed to react overnight. After immobilizing cortisol antibody, the sensor was rinsed with phosphate-buffered saline (PBS; pH=7.4) and blocked with Bovine serum albumin (BSA, 1%) to prevent non-specific interaction with the immunosensor. The developed cortisol immunosensor was stored at -20 °C until further use.

### 2.3. Characterization of developed multilayer immunosensor for cortisol detection

The developed sensor was characterized during each fabrication step to carefully evaluate the effect of different coating and functionalization steps using various characterization techniques. Atomic force microscopy (AFM, MFP3D-BIO, Asylum Research) study was carried out to validate the thickness (by scratch method) and roughness profile of different coating layers during sensor fabrication. Attenuated total reflectance-Fourier transform infra-red (ATR-FTIR, NICOLET-iS50 spectrometer, Thermo Fischer) was performed to analyze surface functional groups, validate functionalization, and conjugation of biorecognition (antibody/protein) element on the sensor surface. X-ray photoelectron spectroscopy (XPS, AXIS Supra^+^, Kratos Analytical) was used to analyze the elemental composition of different coated layers. Change in the surface chemistry of the sensor during each fabrication step was also measured indirectly using water contact angle measurement.

To highlight and provide proof of concept for conjugation of antibodies using EDC-NHS coupling. We used fluorescence-tagged fibrinogen protein (Alexa fluor 647 fibrinogen) to immobilize the sensor surface using EDC-NHS coupling as per the above protocol used for antibody conjugation. In brief, the EDC-NHS mixture was treated with Alexa fluor 647 fibrinogen at a concentration of 1 mg/mL. The EDC-NHS-activated fibrinogen solution was added to the sensor surface and allowed to react overnight. After conjugation, the sensor was rinsed with PBS solution. The presence of conjugated fluorophore protein on the sensor surface was imaged using a fluorescence microscope (ZEISS Axio Imager 2) under a red emission channel (excitation 650 nm and emission 680 nm).

### 2.4. Detecting Cortisol using electrochemical sensing

Cortisol stock solution (Sigma Aldrich, cat. #H4001) was prepared by dissolving 1 mg of hydrocortisone in 100 μL of ethanol and then diluted it with 900 μL of PBS buffer to achieve a final concentration of 1 mg/mL as per supplier protocol. The prepared cortisol stock solution was further diluted to form 1 μg/mL of working solution. The sensor’s impedance was recorded using a Bode 100 impedance analyzer (Omicron Lab). At first stable connection on the glass substrate was secured using a customized in lab developed 3-D printed platform embedded with pogo pins. To obtain the impedance characteristics of the fabricated sensor, a two-port electrical impedance measurement technique was used. A sinusoidal excitation voltage of 13 dBm was applied to the input port. The excitation frequency was logarithmically swept from 100 Hz to 40 MHz in 200 steps. Initially, the impedance reading of the sensor was measured after each coating and functionalization layer. Later, the change in impedance upon adding cortisol was recorded with samples spiked with 50, 100, and 200 ng/mL of cortisol. To assess the specific binding of cortisol with antibodies, the impedance reading of the sensor was also measured using a blank PBS solution and negative control LDL (Low-density lipoprotein). All the electrochemical sensing tests were carried out in multiple replicates (n = 5), and average values were represented.

### 2.5. Statistical analysis

A one-way ANOVA test was performed to assess statistically significant differences in impedance value among three layers and cortisol dose response. A p-value less than 0.05 was considered statistically significant and represented by respective symbols.

## 3. Results and Discussion

### 3.1 Multilayer immunosensor development

Development of non-Faradic EIS requires a conductive substrate immobilized with biomolecules (like antibodies or aptamers) to evaluate change in its electrical properties (capacitance or impedance or resistance) upon interaction with tested bio-analyte [32]. To develop a conductive immunosensor platform, we spin-coated one of the most commonly used conductive organic polymers (PEDOT: PSS) on clean etched ITO-coated glass [33]. The obtained PEDOT: PSS (ITO-PS) surface lacks the chemical functional groups needed to conjugate biomolecules. Therefore, we followed a classical approach of adding a linker molecule on the PEDOT: PSS surface to conjugate antibodies. ATPES is one such standard linker molecule, which has been frequently used to conjugate biomolecules like proteins and antibodies on the PEDOT: PSS surface [31,34,35]. To couple APTES on PEDOT: PSS surface, spin-coated PEDOT: PSS film was treated with UV-Ozone to impart hydroxyl functional groups on the film surface, enabling intermediate APTES coupling through silanization [36]. Obtained APTES-coated ITO-PS substrate (ITO-PS-A) has linker APTES that provides the free amine group on the substrate (ITO-PS-A) for protein/antibodies conjugation through a well-established EDC-NHS chemistry [30]. Using EDC-NHS coupling reaction, we conjugated cortisol antibody on (ITO-PS-A) surface using carbamide reaction between the free amine group of APTES and the carboxyl group of antibody to develop cortisol immunosensor (ITO-PS-A-Ab) as illustrated in **Fig. 2**.

**Fig. 2:**
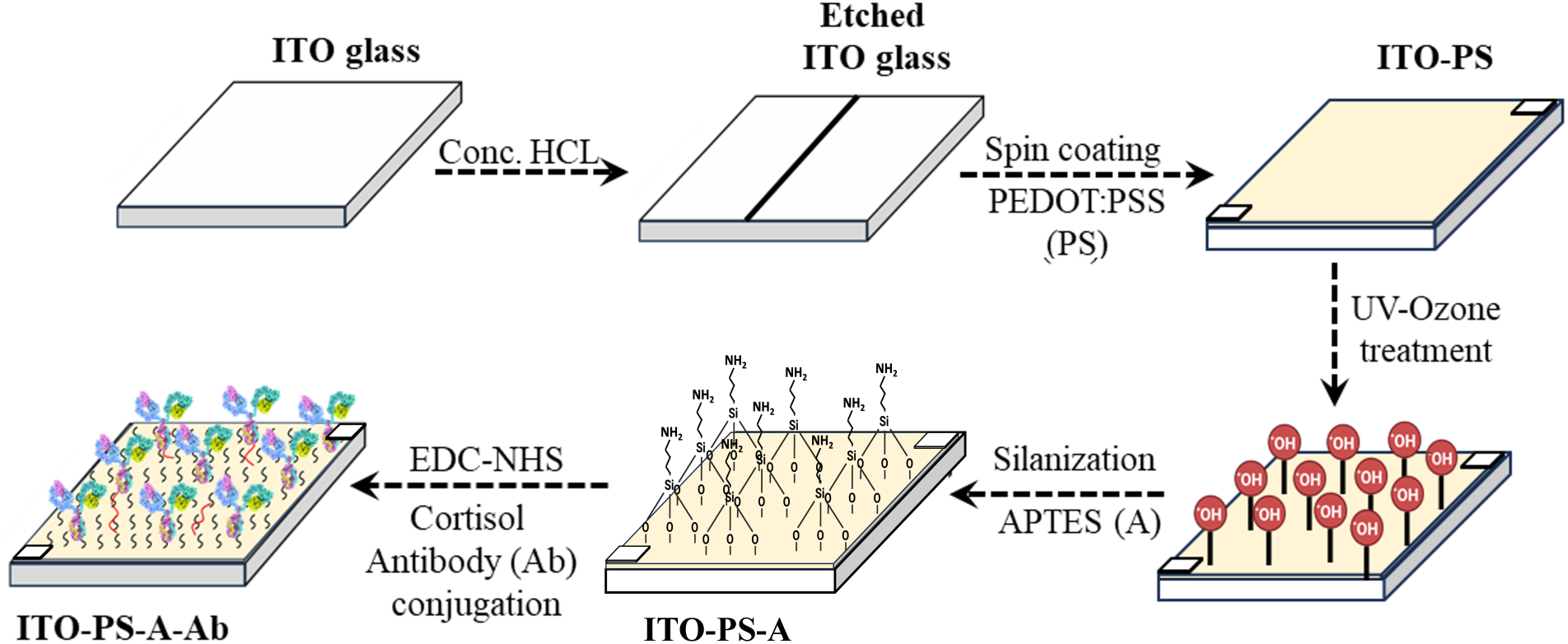
Schematic illustration of multilayer sensor fabrication by sequential coating and immobilization of PEDOT: PSS, APTES, and antibody on ITO glass substrate.

### 3.2. Physicochemical characterization of multilayer immunosensor

Fabricated cortisol immunosensor by the subsequent layering of different components on ITO glass was thoroughly characterized to confirm the presence, morphology, and chemical characteristics of different coated/conjugated layers. Coated PEDOT: PSS film showed uniform coverage on ITO-coated glass with a thickness of around 400 nm. Coupling of APTES on PEDOT: PSS surface resulted in an increase in the film thickness from 400 nm to 700 nm **(Fig. 3(i))** and roughness (Ra) from 1.7 nm to 9.74 nm **(Fig. 3(ii)),** suggesting coating of APTES on PEDOT: PSS surface. Chemical conjugation of APTES and antibody on PEDOT: PSS surface was confirmed using FTIR, shown in **Fig. 3(iii).** FTIR spectra of APTES coated PEDOT: PSS film showed characteristic peak for PEDOT: PSS (C=C stretching of cyclic alkene at 1705 cm^-1^ and S=O stretching of sulfate group at 1290 cm^-1^ [37,38]) and APTES (Si-O-C stretching at 950 cm^-1^ [41]). In addition to these peaks, APTES-coated PEDOT: PSS film showed an additional peak at 1108 cm^-1^ corresponding to Si-O-Si stretching, confirming the conjugation of the APTES layer on PEDOT: PSS [39]. Antibody-coated PEDOT: PSS-APTES sampled showed broad OH stretching peaking (3450 cm^-1^) and broad amide I (1640) peaks confirming bioconjugation of cortisol antibody on APTES surface for specific binding of cortisol [40].

**Fig. 3:**
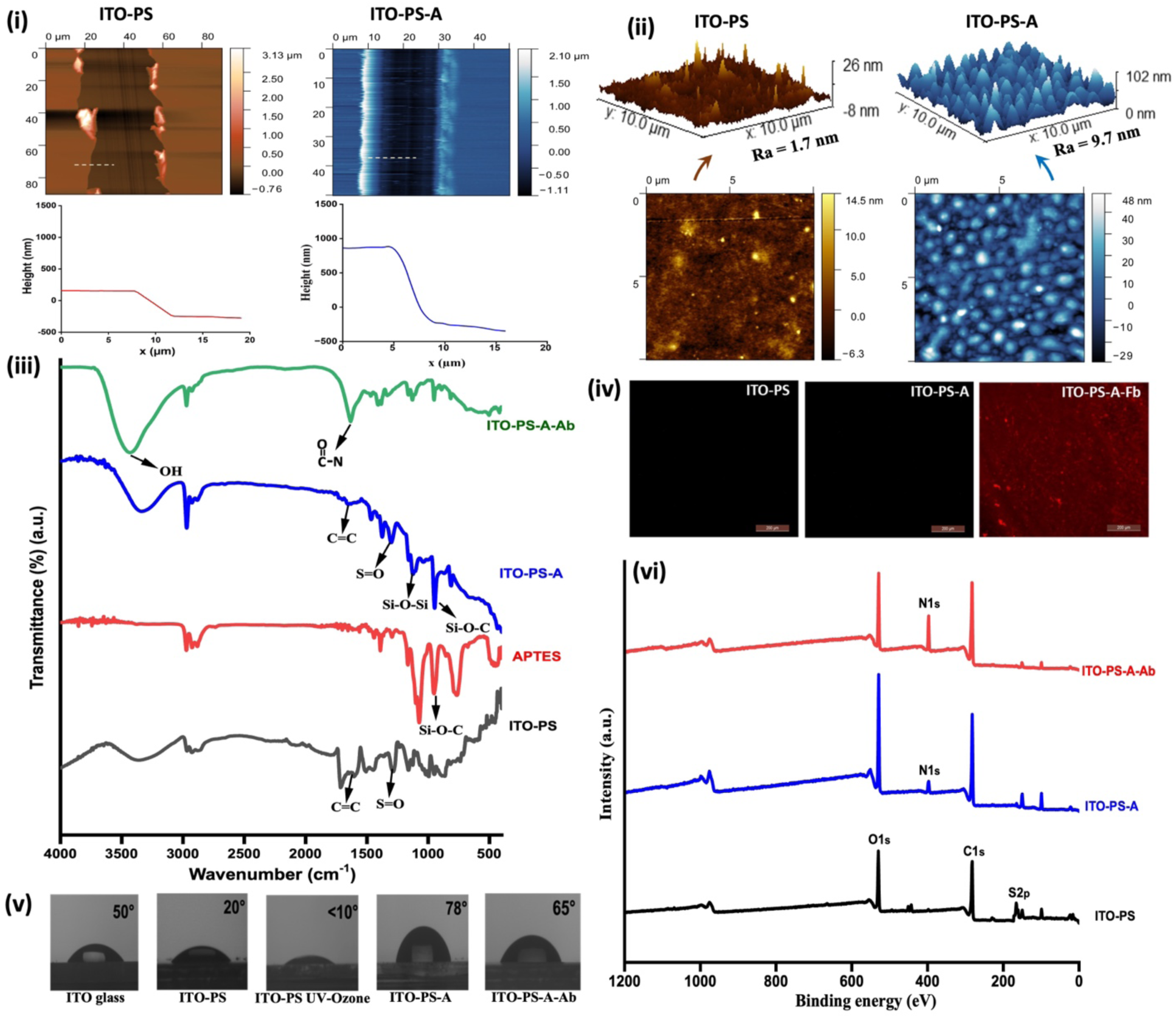
Physicochemical characterization of developed multilayer immunosensor. (i) and (ii) AFM thickness and roughness profile of PEDOT: PSS (ITO-PS) and APTES (ITO-PS-A), (iii) FTIR spectra of PEDOT: PSS, APTES, and antibody (ITO-PS-A-Ab) immobilized on the sensor surface, (iv) Fluorescent micrograph of PEDOT: PSS, APTES, and fluorophore Fibrinogen (ITO-PS-A-Fb) coated sensor confirming the presence of fluorescent protein after EDC-NHS coupling (scale bar = 200 µm), (v) Contact angle of the sensor surface during different fabrication steps, (vi) XPS spectra of PEDOT:PSS, APTES and antibody layer.

To provide proof of concept for conjugation of antibodies using EDC-NHS coupling. We used fluorescence-tagged fibrinogen (Fb) protein to immobilize the sensor surface. Fluorescence image of PEDOT: PSS and PEDOT: PSS-APTES surface showed no fluorescence background in the red channel. After fluorophore fibrinogen conjugation on the ITO-PS-A substrate, it showed red fluorescence spots all over the sensor surface (ITO-PS-A-Fb), indicating uniform immobilization of protein. It provided a clear picture and proof of concept that the conjugation of antibodies using EDC-NHS would be very similar to that of fibrinogen, shown in **Fig. 3(iv).**

Chemical conjugation of different molecules on a sensor changes the surface’s chemical properties. The Change in surface chemistry of PEDOT: PSS film upon APTES and antibody coating was evaluated using XPS (**Fig. 3(v)**). XPS spectra of spin-coated PEDOT: PSS showed the presence of elemental peaks for carbon (C1s, 282 eV), oxygen (O1s, 530 eV), and doublet Sulphur (2Ps, 165 eV and 161 eV), confirming the presence of organic film with presence of sulfur groups originating from PEDOT: PSS coating on the glass substrate [41]. On the other hand, APTES-coated PEDOT: PSS film showed an additional peak for Nitrogen (N1s) at 398 eV corresponding to the free amine group from APTES molecules, confirming the presence of APTES on PEDOT: PSS surface for linking antibody [42,43]. ITO-PS-A-Ab substrate showed a further increase in elemental Nitrogen (N1s) peak intensity indicating the presence of nitrogen-rich biomolecule (antibody). Overall using XPS, we confirmed the change in surface composition upon conjugation of APTES and antibody on the PEDOT: PSS surface. Change in the surface chemistry of the developed sensor during subsequent coating was also confirmed using water contact angle measurement (**Fig. 3(v&vi)**). Clean ITO glass showed water contact angle of approximately 50°, which decreased to 20° after coating of hydrophilic PEDOT: PSS polymer film. UV-Ozone treatment of PEDOT: PSS film imparted polar hydroxyl group on the surface, which further lowered the contact angle to less than 10°. Upon conjugation of hydrophobic APTES on the PEDOT: PSS surface, the water contact angle increased to 78°. Finally, immobilization of polar-natured antibodies results in a decrease in water contact angle to 65°. These sequential changes in water contact angle further substantiate subsequent modification and the multilayer nature of the developed immunosensor.

### 3.3. Electrical characterization of multilayer immunosensor

The electrical characteristics study of the developed sensor was inspected by non-faradic EIS to determine the change in impedance after the binding of the analyte (cortisol) with the biorecognition element (cortisol antibody). Non-faradic EIS provides valuable insights into the impedance response (capacitance and resistance) of the double layer formed at the electrode surface [44]. To measure the impedance change of the developed sensor, we developed an in-lab 3D-printed assembly having metallic pogo pins to ensure the proper connection and repeatability in measurement with the electrode surface on the sensor substrate. An impedance test fixture was used to obtain precise measurements during the subsequent development of the multilayer sensor platform shown in **Fig. 4 ((i) and (ii))**. After calibration, the impedance of individual coating on ITO glass was recorded in the frequency range of 100 Hz to 40 MHz. EIS data demonstrated that the change in the impedance values was found to be consistent between 1 kHz and 10 MHz. Therefore, all the impedance readings on the sensor substrate were measured at 10 kHz and highlighted in **Fig. 4(iii)**. The coating of conductive polymer (PEDOT: PSS) on ITO glass showed an average impedance of around 259 Ω. Following APTES coating on top of PEDOT: PSS sensor surface, the impedance of ITO-PS-A film increased to 630 Ω attributing to the non-conductive nature of APTES. To confirm the substantial rise in the impedance was due to APTES’s non-conducting behavior instead of increased thickness, we prepared a spin-coated PEDOT: PSS layer of thickness approximately 700 nm (comparable to APTES/PEDOT: PSS layer) on the ITO glass substrate and its impedance was measured. The impedance of 700 nm thick PEDOT: PSS film was found to be 310 Ω which was drastically different from ITO-PS-A film, confirming that the considerable impedance difference of APTES coated PEDOT: PSS film was attributable to APTES’s non-conducting nature. Later, antibodies were immobilized on PEDOT: PSS-APTES sensor surface using EDC-NHS chemistry, which further increased the impedance value to 684 Ω. The increase in impedance upon antibody bioconjugation is indicative of the introduction of an insulating layer on the electrode surface [45].

**Fig. 4:**
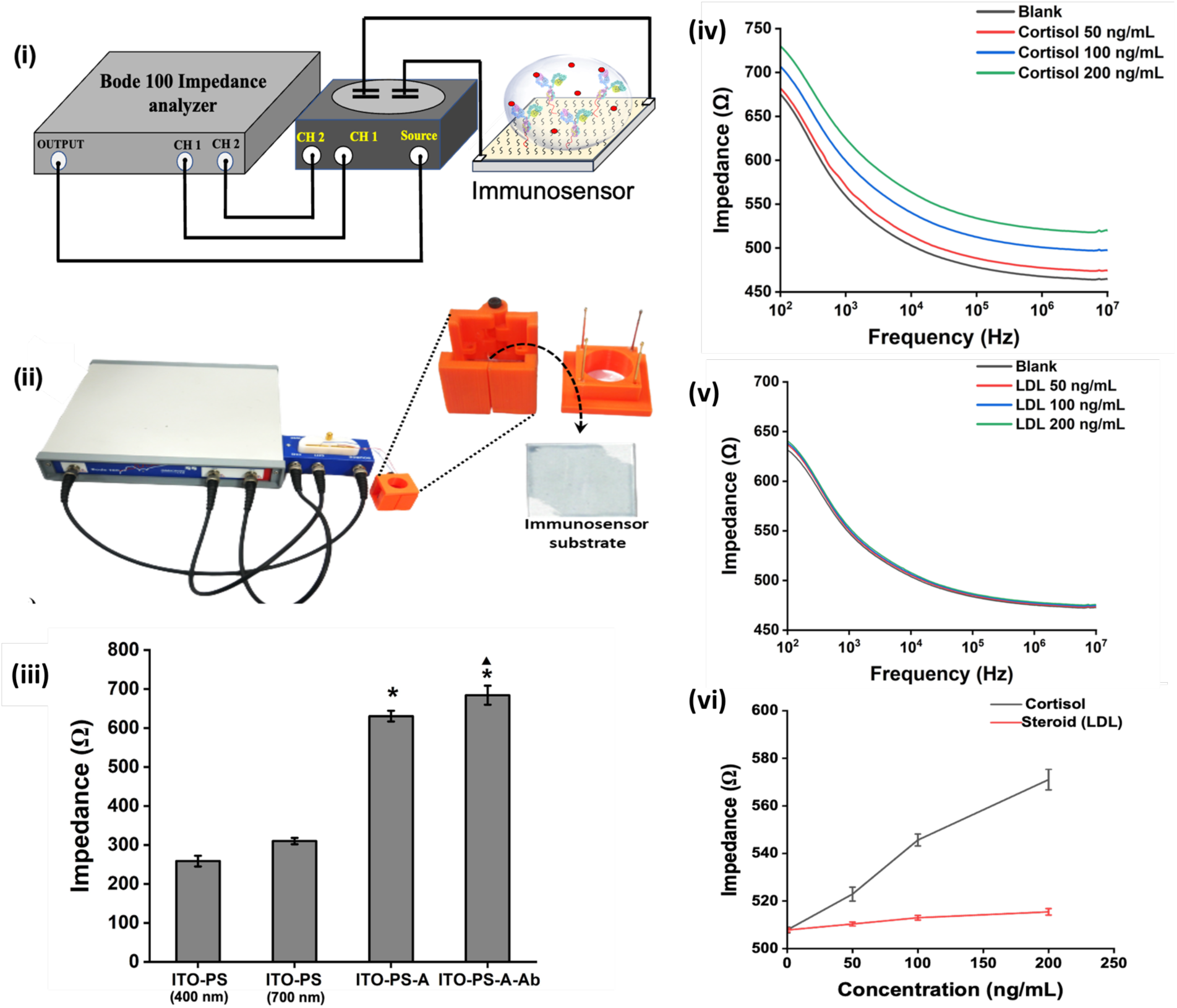
Impedance response of developed immunosensor. (i) Schematic illustration of sensor assembly for impedance reading, (ii) Assembly of developed cortisol immunosensor and 3-D printed customized platform for impedance measurement using Bode 100 impedance analyzer, (iii) Impedance reading of PEDOT: PSS, APTES, and antibody functionalized sensor surface at 10 kHz; Statistically significant difference (p < 0.05) compared to ITO-PS and ITO-PA-A are indicated by * and ▴ respectively, (iv) and (v) Concentration dependent bode plot for cortisol and LDL at a frequency range of 100 Hz to 40 MHz, (v) Comparative concentration dependent bode plot for cortisol and LDL at a frequency of 10 kHz

Finally, cortisol in PBS (analyte) was added on PEDOT: PSS-APTES-Ab surface (50-200 ng/mL) to evaluate the impedance change upon binding with cortisol antibody. **Fig. 4(iv)** shows cortisol dose-dependent change in impedance. The binding of cortisol with antibodies has been shown to block the flow of electrons to the sensor surface (acting as an additional insulating layer), which increases impedance with the increase in cortisol concentration [44]. To confirm the specific binding of cortisol with antibodies and prevent electron flow. We added a non-specific steroid-based analyte LDL in the range of 50 -200 ng/ml on PEDOT: PSS-APTES-Ab surface and measured the impedance (**Fig. 4(v)**). Adding non-specific LDL analyte on PEDOT: PSS-APTES-Ab surface showed negligible change in impedance with the increase in LDL concentration in comparison to cortisol **(Fig. 4(vi)),** suggesting the flow of electrons from the electrolyte solution to the electrode surface remains unimpeded. Since LDL molecules do not have binding affinity for the antibodies, they may not interact with the sensor surface. As a result, there will be no change in electron flow. These results indicated that our developed sensor displayed good selectivity towards cortisol without interfering with non-specific LDL molecules.

### 3.4. Equivalent circuit modelling and mechanism

The obtained impedance data of cortisol collected from the developed multilayer immunosensor was fitted through an equivalent circuit model using ZSimpWin software, shown in **Fig. 5(i – iii)**. The model comprises various components that contribute to its overall behaviour, which includes bulk solution resistance (Rs), charge transfer resistance (Rct), and capacitance (C1), representing resistance encountered by ions flowing through the solution and cortisol-antibody interaction. Furthermore, the model incorporates constant phase element, resistance, and capacitance of ITO-PS-A electrodes, which play a crucial role in the overall functioning. To minimize the influence of Warburg impedance, all readings were taken at higher frequencies (100 Hz – 40 MHz) [46]. As the cortisol dose concentration increases, there is a corresponding increase in cortisol-antibody complexes which leads to a change in the electrical properties of the electrode surface. Alteration in electrical properties results in the formation of an additional insulating layer on the sensor surface which impedes the electron flow from the solution to the electrode surface. The reduced flow of electrons corresponds to an increase in the impedance (Zreal and -Zimg) of the overall system, demonstrated by the Nyquist plot (**Fig. 5(iv)**) and illustrated in **Fig. 5(v)**. The increase in real and imaginary impedance was further demonstrated by the increase in Rct and decrease in C1, respectively. The theoretical fit of the experimental results of the developed biosensor using resistance, capacitance, and constant phase element are summarized in Table 1, which shows an increase in Rct and a decrease in C1 with the rise in cortisol concentration. The equivalent model of cortisol dose response was mostly dominated by two components (Rct and C1) and indicated that the sensor is detecting resistive change resulting from the specific molecular binding of the cortisol to the antibodies. On the other hand, when a solution with a non-specific analyte is introduced, it does not interact at the molecular level, and hence it does not bind with cortisol-antibodies present on the sensor surface. As a result, the current flow at the electrode surface was not altered, as reflected in the overall impedance of the sensor.

**Fig. 5:**
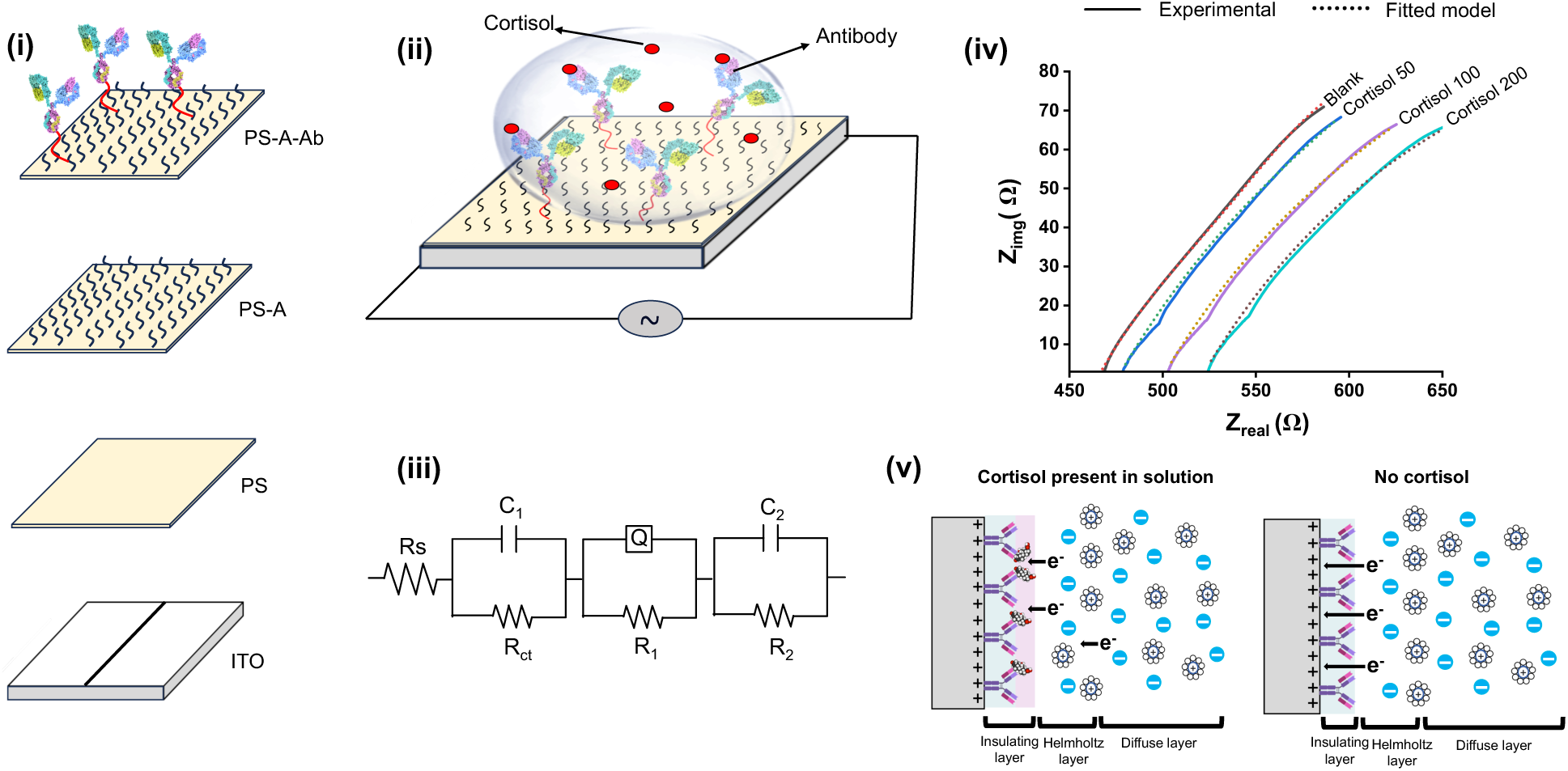
Equivalent circuit of multilayered immunosensor and its working mechanism. (i) Schematic representation of multilayer cortisol biosensor, (ii) Diagrammatic representation of cortisol detection on the sensor surface using antibodies as a biorecognition element, (iii) Equivalent circuit model of developed sensor, (iv) Nyquist plots from the EIS measurements of developed sesnor at increasing cortisol concentration, (v) Schematic showing increased thickness of insulating layer after binding of cortisol with antibodies on the sensor surface

**Table 1:** Theoretical fit of experimental results for constant phase element, capacitive and resistive components of equivalent circuit model.

| **Cortisol**<br>**concentrat**<br>**ion**<br>**(ng/mL)** | **Components of equivalent circuit model** |  |  |  |  |  |  |
| --- | --- | --- | --- | --- | --- | --- | --- |
|  | **R<sub>s</sub>** | **R<sub>ct</sub>** | **C<sub>1</sub>** | **R<sub>1</sub>** | **Q** | **R<sub>2</sub>** | **C<sub>2</sub>** |
| **50** | 5628<br>±25.773 | 473.6<br>±5.9823 | 0.001328<br>±0.000862 | 3550<br>±83.219 | 0.0003023<br>±0.0001034 | 483.6<br>±15.324 | 0.002955<br>±0.00198 |
| **100** | 5908<br>±32.556 | 497.4<br>±9.324 | 0.001099<br>±0.000435 | 714.8<br>±109.81 | 0.000274<br>±0.000912 | 428.8<br>±24.764 | 0.002173<br>±0.00104 |
| **200** | 5219<br>±17.314 | 520.5<br>±7.723 | 0.0006036<br>±0.0000542 | 2403<br>±60.527 | 0.0002462<br>±0.000634 | 380<br>±34.438 | 0.0009777<br>±0.000537 |

## 4. Conclusions

Multilayer non-faradic immunosensor was developed for the specific detection of cortisol. The multilayer immunosensor was fabricated by coating a PEDOT: PSS on ITO glass followed by anti-cortisol antibody immobilization on the surface through a linker APTES molecule. The coating of different layers resulted in a change in the morphology and physicochemical properties of the substrate. AFM micrograph showed an increase in thickness and roughness profile upon coating PEDOT: PSS and ATPES on ITO glass substrate. FTIR, XPS, and water contact measurement confirmed coating and immobilization of PEDOT: PSS, ATPES, and cortisol antibody to form an immunosensor for cortisol detection. The electrical characteristic of different coating layers showed a change in impedance due to the formation of subsequent resistive layers. The developed immunosensor showed real-time specific detection of cortisol in a physiologically relevant range of 50-200 ng/mL.

## Ethics approval and consent to participate

Not applicable

## Availability of data and materials

The datasets generated during and/or analyzed during the current study are available from the corresponding author on reasonable request

## Acknowledgments

The authors would like to acknowledge the IIT Delhi core research facility (CRF) and no research facility (NRF) for characterization. The authors would like to thank Anne Tryphosa for assistance in impedance test. We would like to thank Maneesha Tewari for helping in protein conjugation. We would also like to thank Shivani Chaudhary and Doyel Ghosal for insightful discussions and sharing expertise on material characterization and resources.

## Contribution

C.G., S.K., and B.M. jointly conceptualize the work, C.G. conducted the research work, compiled data, and wrote the manuscript. S.K. and B.M. analyzed that data and edited the manuscript. S.K.P provided technical input on sensor design and edited the manuscript.

## Vitae

**Chinmay Gupta** is a master’s student at Centre for Biomedical Engineering, Indian Institute of Technology Delhi, New Delhi, India. He completed his undergrad in Veterinary medicine from G.B. Pant University of Agriculture and Technology, US Nagar, Uttarakhand, India. His area of research is Biosensors, Point-of-care diagnostics devices, Microfluidics, and 3-D printing.

**Sudip Kumar Pattanayek** is a Professor at Department of Chemical Engineering, Indian Institute of Technology Delhi, New Delhi, India. His area of research is structure and dynamics of Proteins at Interfaces, Aggregation of Proteins, Polymer-surfactant interactions, Rheology of dilute polymer solution, Polymer nanocomposites, Interfacial Rheology, Rheology of slurry.

**Biswarup Mukherjee** is an Assistant Professor at Centre for Biomedical Engineering, Indian Institute of Technology Delhi, New Delhi, India. His area of research is focused towards developing multi-modal platform technologies that provide the means to non-invasively monitor aspects of neuromuscular activity in real-time, particularly for rehabilitation applications. Development of biomimetic sensors to improve sensorimotor integration in individuals with motor disabilities.

**Sachin Kumar** is an Assistant Professor at Centre for Biomedical Engineering, Indian Institute of Technology Delhi, New Delhi, India. His area of research underlies in interdisciplinary field of biomaterials, biomechanics, and bio-optics to understand how mechanical, chemical, and physical properties of the biomaterials regulate cellular functions and processes such as cancer progression, stem cell differentiation, tissue regeneration and cellular metabolism.

